# Replication of early and recent Zika virus isolates throughout mouse brain development

**DOI:** 10.1101/177816

**Authors:** Amy B Rosenfeld, David J Doobin, Audrey L Warren, Vincent R Racaniello, Richard B Vallee

## Abstract

Fetal infection with Zika virus (ZIKV) can lead to congenital Zika virus syndrome (cZVS), which includes cortical malformations and microcephaly. The aspects of cortical development that are affected during virus infection are unknown. Using organotypic brain slice cultures generated from embryonic mice of various ages, sites of ZIKV replication including the neocortical proliferative zone and radial columns, as well as the developing midbrain, were identified. The infected radial units are surrounded by uninfected cells undergoing apoptosis, suggesting that programmed cell death may limit viral dissemination in the brain and may constrain virus associated injury. Therefore, a critical aspect of ZIKV induced neuropathology may be defined by death of uninfected cells. All ZIKV isolates assayed replicated efficiently in early and mid-gestation cultures, and two isolates examined replicated in late-gestation tissue. Alteration of neocortical cytoarchitecture, such as disruption of the highly-elongated basal processes of the radial glial progenitor cells, and impairment of postmitotic neuronal migration were also observed. These data suggest that all lineages of ZIKV tested are neurotropic, and that ZIKV infection interferes with multiple aspects of neurodevelopment that contribute to the complexity of cZVS.

**Significance:** Zika virus infection has been associated with multiple pathologies of the central nervous system (CNS) including microcephaly, Guillain-Barré syndrome, lissencephaly, the loss of white and grey matter volume and acute myelitis. Using organotypic brain slice cultures, we determined that ZIKV replicates across different embryonic developmental stages, and viral infection can disrupt proper brain development leading to congenital CNS complications. These data illustrate that all lineages of ZIKV tested are neurotropic, and that infection may disrupt neuronal migration during brain development. The results expand our understanding of neuropathologies associated with congenital Zika virus syndrome.

## Introduction

Microcephaly is a neurodevelopmental disorder that is clinically characterized by dramatic physical reduction of head circumference (1-3). A variety of etiologies can lead to microcephaly, including vertical transmission of the TORCH microbes: ***T****oxoplasma gondii*, **R**ubella, **C**ytomegalovirus, **H**erpes simplex virus, or **O**ther pathogens, such as Coxsackievirus, varicella zoster virus, HIV, human T-lymphotropic virus, or *Treponema*. Infection with the arbovirus ZIKV has been associated with the development of congenital cortical malformations including microcephaly (4-15), defining the virus as a new TORCH agent (10, 16).

The reduced head circumference seen with microcephaly is a consequence of reduced brain volume, also manifested in a thinner neocortex. A healthy mature cerebral cortex is composed of six distinct layers of neurons. Most cells within these layers arise from a single layer of apical-basal polarized cells that line the cerebral ventricle (17-23). This single layer of cells undergoes multiple rounds of symmetric mitotic divisions, which largely define the ultimate size of the brain. These cells increasingly undergo asymmetric divisions at which point the cells are referred to as radial glial progenitors (RGPs) (24-26). The asymmetric divisions create two postmitotic daughter cells, with one migrating basally out from the proliferative ventricular zone (VZ) towards the developing cortical plate (CP) of the neocortex. The other cell remains as an RGP attached at the ventricular surface, and the collective progeny of this RGP defines a “radial column.” A signature feature of the RGP is its elongated basal fiber that links the VZ to the opposing pial surface. Migration of the postmitotic cells into cortical plate of the developing neocortex occurs along the RGP process (24, 27-31). The late stages of neocortical development establish the gyri, sulci, and cytoarchitecture of the mature brain. Mutations in genes critical for these processes all may interfere with the proper number of neurons and layers, resulting in a thinner neocortex, consistent with autosomal recessive microcephaly (reviewed in (32)).

Zika virus, a (+) RNA arbovirus in the *Flaviviridae* family, is a re-emerging human pathogen. Originally isolated from a febrile sentinel macaque in 1947, the first reported human infections occurred in Nigeria during the early 1950s (33-36). Initially it was thought that Zika virus infection did not result in clinical disease, but was experimentally neurotropic in mice (33, 36). If symptoms of viral infection were observed in humans, they were mild, including rash, joint pain, and fever (37-43). During the 2007 and 2013 outbreaks in isolated Pacific islands, adult ZIKV infection was associated with neurological dysfunction such as Guillain-Barré syndrome, a disorder in which the immune system attacks the peripheral nervous system, and other central nervous system (CNS) complications (40, 44-54). Once ZIKV spread to the Western Hemisphere, it was recognized to be responsible for the increase of children born with microcephaly and cortical malformations in Brazil and Colombia (9, 55-59). By late 2016, ZIKV infection was known to cause many neurological abnormalities outside of microcephaly, including lissencephaly, loss of brain volume, pachygria, and ventriculomegaly (8, 11, 12, 60-64), which all comprise congenital Zika virus syndrome (cZVS) (4, 5, 8, 11-15).

Several animal models of ZIKV infection including mice and non-human primates have recently been established (65-88). These models illustrate many features of ZIKV induced pathology including viremia and neuronal tissue tropism. The inability to genetically manipulate non-human primates restricts their use for understanding cZVS. Organoid culture models provide limited information about virus infection of target cells within the context of the developing tissue and organ, since some cell types are missing and there is little to no immune system or vascular representation. Furthermore, many organoid cultures used to understand ZIKV biology only develop neural identity, not tissue organization (89, 90). Organoid cultures do not allow for the study of neuropathologies in the late developing brain, are difficult to consistently manufacture, and are characterized by immature neuronal connectivity; therefore, they are far more simplistic than the brain (91).

Organotypic brain slice cultures are widely used to understand neuronal connectivity and neurodegenerative disorders (92-94), and have enriched our understanding of autosomal-recessive microcephaly and brain development (95, 96). Furthermore, organotypic brain slice cultures from embryonic mice are a genetically amenable system to study cZVS and the ability of the virus to infect neural cells (neurotropism), as the virus inoculum is placed directly on target cells of the cultures. This system fully separates neurotropism from neuroinvasion, virus entry into the CNS from distant sites of the body. Consequently, observations made in organotypic brain slice cultures generated from embryonic mice expand our understanding of viral infection beyond established animal models and organoid cultures.

In this study the effects of ZIKV infection on the embryonic mouse brain were determined using organotypic brain slice cultures. All ZIKV isolates tested from 1947 to the present replicated in early and mid-gestation embryonic brain tissue (E13 and E15, respectively), while two isolates replicated in brain slice cultures from E19 embryos. Virus replication lead to increased apoptosis within the cultures and impaired neuronal migration into the cortical plate. The results suggest that infection with different isolates of ZIKV can cause severe brain developmental abnormalities. It is likely that brain complexity is reduced through multiple mechanisms during ZIKV infection. These data begin to illuminate how ZIKV-associated neuropathologies develop.

## Results

### Zika virus neurovirulence is independent of genetic lineage

It has been suggested that neurovirulence was acquired as ZIKV spread from Africa though Asia and Pacific to South America (97). Analysis of the genome sequences of isolates from 1947 to the present defines two phylogenetic groups (Fig. 1). Neurologic disease has been associated with the Asian lineage of viruses (97). To determine if neurotropism is a recently acquired property, replication of virus isolates from both phylogenetic groups was assayed in organotypic brain slice cultures from E15 mouse embryos. Neurogenesis at this stage corresponds to weeks 13/14 in human gestation, the beginning of the second trimester when there is maximal migration of the post-mitotic neurons into the cortical plate (21, 98). Indirect immunofluorescence microscopy (IF) of brain slice cultures 3 days post infection (dpi) with the 2015 Puerto Rican ZIKV isolate revealed virus specific antigen (ZIKV-E) throughout the developing neocortex and midbrain (Fig. 2A). Notably, ZIKV-E was visualized in cells that line the ventricular surface of the developing brain, with staining extending throughout the radial columns of progenitor cells and their neuronal progeny (Fig. 2B). This observation aligns with previous reports that human and mouse neuronal stem cells are a site of ZIKV infection (99, 100). Furthermore, the same phenotype, namely the presence of viral antigen present throughout the cortical plate to the pial surface, was observed in brain slices infected with an isolate acquired from Malaysia during 1996 (Fig. S1).

**Figure 1.**
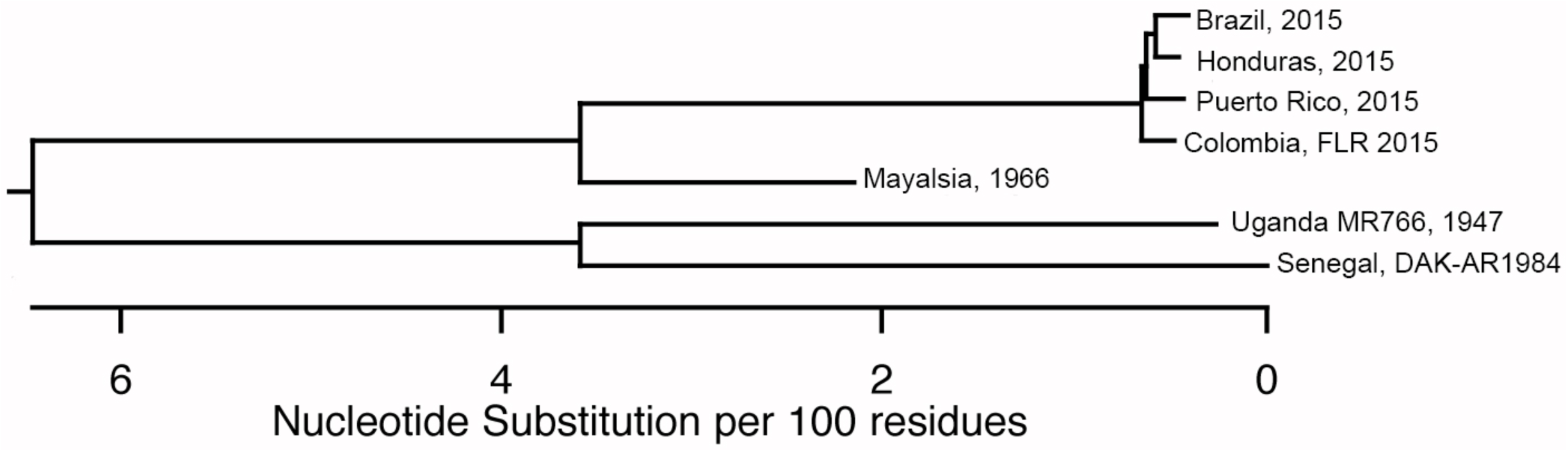
Phylogenetic tree of ZIKV isolates used in this work. Entire genome sequences of 7 ZIKV isolates examined in this study were aligned using the ClustalW module of DNAstar.

**Figure 2.**
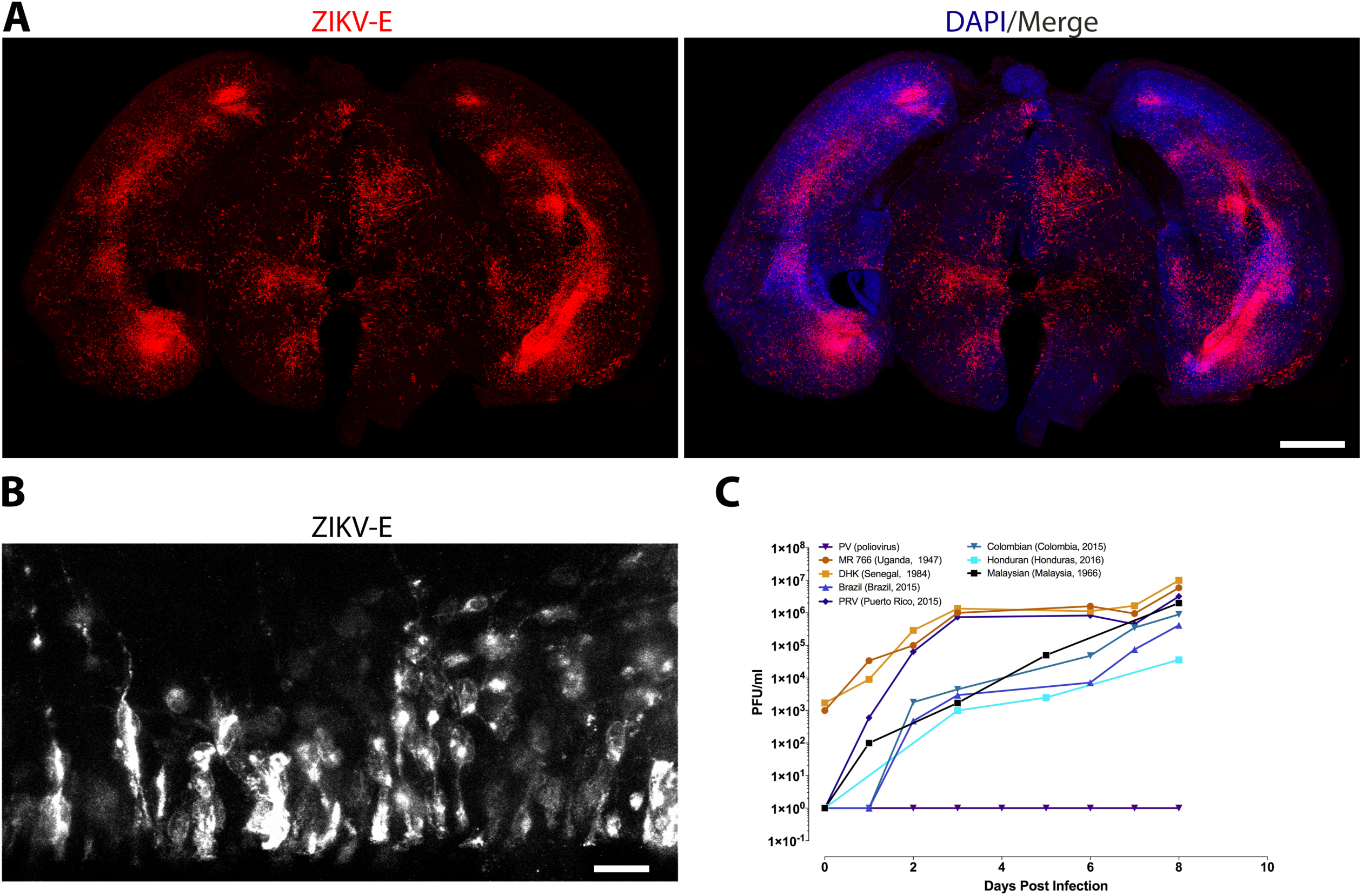
Representative tissue distribution and replication of ZIKV isolates in organotypic mouse brain slices. **(A)** Brain slice cultures from E15 embryos were infected with 10^5^ pfu of the ZIKV PRV isolate, and at 4 dpi were fixed and stained with pan-flavivirus antibody against the E glycoprotein (ZIKV-E) and DAPI. Scale bar is 500 µm. **(B)** Higher magnification (60x objective) of the ventricular surface of infected brain slice culture, stained with pan-flavivirus antibody. Scale bar is 10 µm. **(C)** Time course of replication of multiple isolates of ZIKV in E15 brain slice cultures.

To determine if E15 brain slice cultures could be productively infected, they were infected with 10^5^ pfu of seven ZIKV isolates from 1947 to the present: African [MR766], Senegal, Brazilian, Colombian, Puerto Rican, Honduran and Malaysian (Fig. 2C). At different times after infection, culture supernatants were harvested and virus production was determined by plaque assay. By 8 dpi, infectious virus was detected in cultures infected with each of the isolates (Fig. 2C). Poliovirus was used as a negative control and did not replicate in brain slice cultures, as the mice used do not produce the poliovirus receptor, CD155 (Fig. 2C).

In previous reports ZIKV infection of human neural progenitor cells produced from induced pluripotent stem cells revealed increased apoptotic cell death (100). Therefore IF analysis was used to detect cleaved caspase 3 in the infected brain slice cultures. There was an increase in apoptosis observed in organotypic slice cultures infected with each ZIKV isolate compared with uninfected cultures (Fig. 3A). Notably, induction of caspase 3 cleavage occurred mostly within the cells that surround the ZIKV infected cell (Fig. 3B), which has also been suggested by the results of other studies (101, 102).

**Figure 3.**
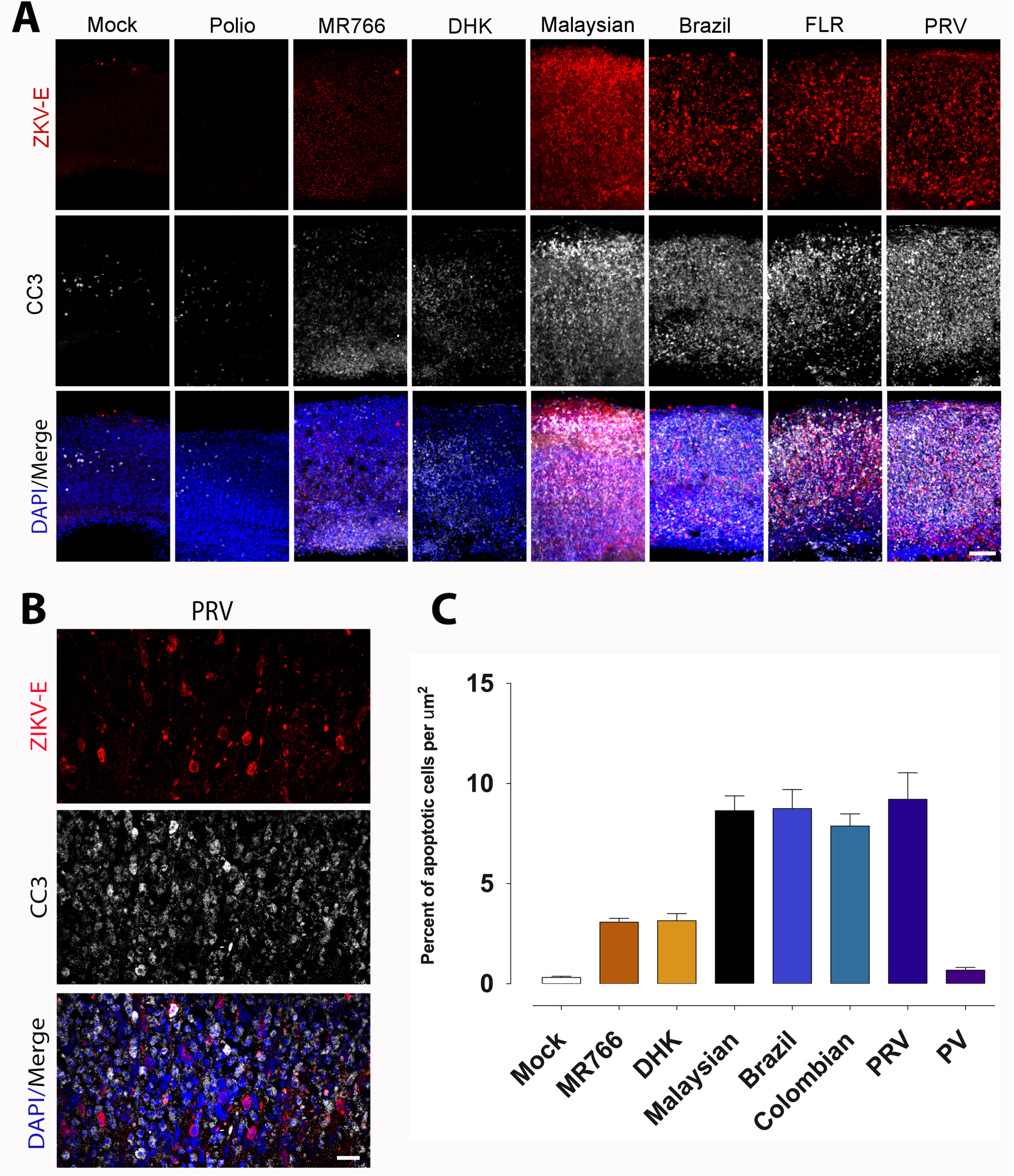
ZIKV infection of organotypic mouse brain slices leads to increased apoptosis. **(A)** Brain slice cultures from E15 embryos were infected with 10^5^ pfu of the indicated ZIKV isolates or poliovirus, and at 8 dpi were fixed and stained with pan-flavivirus antibody against the E glycoprotein (ZIKV-E), antibody to cleaved caspase 3 (CC3), and DAPI. Scale bar is 100 µm. **(B)** Higher magnification (60x objective) of PRV ZIKV infected slices from panel A. Scale bar is 10 µm. **(C)** Apoptotic index, as defined by number of cleaved CC3 positive cells per square micron, in E15 brain slice cultures infected with the indicated ZIKV isolates. PV, poliovirus.

These findings demonstrate that ZIKV infects the embryonic mouse brain during mid-gestation, that the virus is neurotropic independent of its genetic lineage (African or Asian), and that infection leads to apoptosis. The results also establish organotypic embryonic brain slice cultures as a useful system to investigate ZIKV induced brain malformations.

### Developmental restraints of Zika virus infection revealed with embryonic organotypic brain slice cultures

The results of epidemiological studies initially suggested that the greatest risks for developing cZVS are associated with maternal ZIKV infection during the first and second trimester (13, 58). Neurological dysfunction has been associated with virus infection during the 3^rd^ trimester, or shortly after birth, albeit less frequently (103-107). To test whether ZIKV neurotropism is regulated by gestational age, organotypic brain slice cultures were generated from embryonic mice at a range of developmental stages. Brain slices from E13 embryos, which are developmentally comparable to the middle of the first trimester of human neurogenesis, supported viral replication of all 5 ZIKV isolates examined, as determined by expression of a virus specific antigen within focal areas of the neocortex (Fig. 4A, B), and by virus production (Fig. 4C).

**Figure 4.**
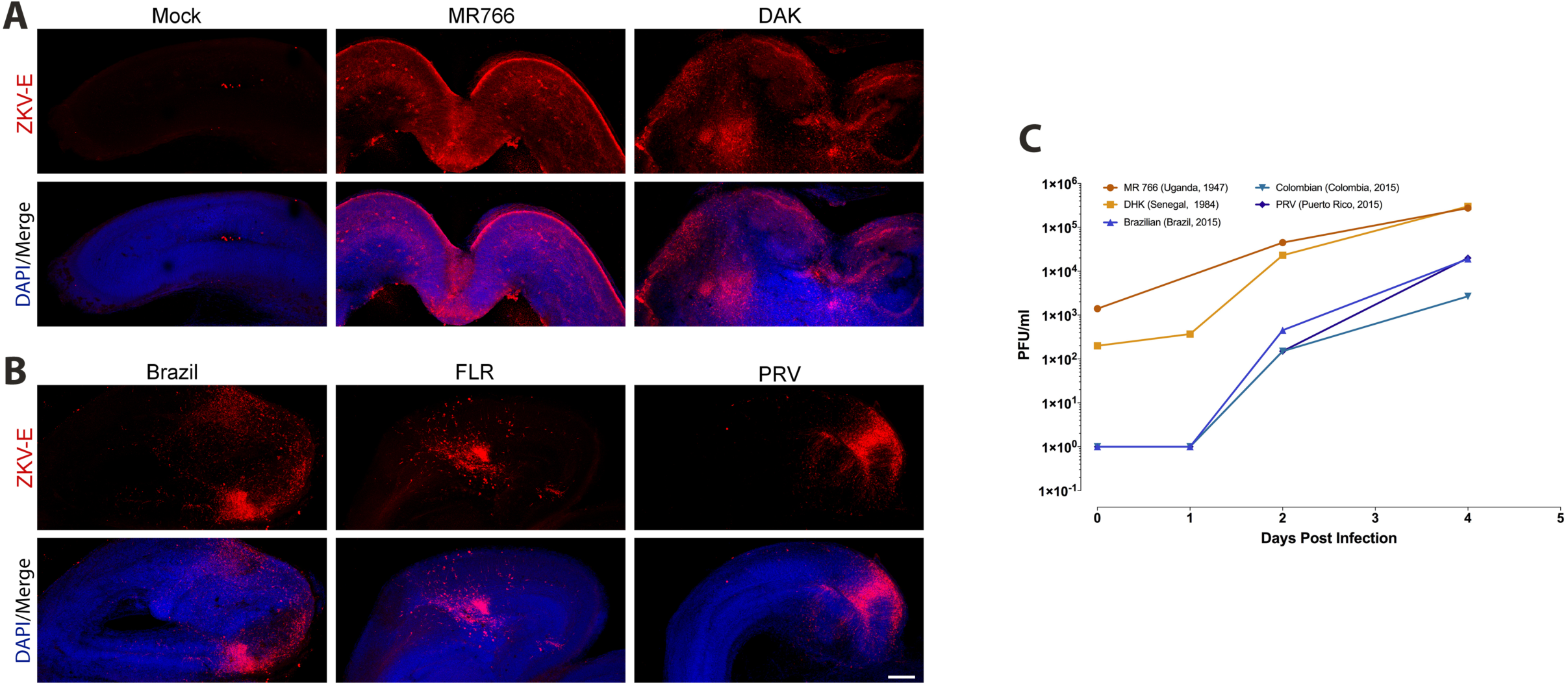
ZIKV replicates in early gestation neural tissue. **(A, B)** Brain slice cultures from E13 embryos were infected with 10^5^ pfu of the indicated ZIKV isolates, and at 8 dpi were fixed and stained with pan-flavivirus antibody against the E glycoprotein (ZIKV-E) and DAPI. Scale bar is 250 µm. **(C)** Time course of replication of multiple isolates of ZIKV in E13 brain slice cultures.

Brain slice cultures derived from E19 embryos, which mirror the late stage of neurogenesis during human gestation, supported productive infection with the Honduran and the prototype MR766 isolates (Fig. 5A). During infection with the Honduran isolate, ZIKV E protein was observed only in the neocortex of E19 slices, not within the proliferative zone as seen in E15 infected slices (Fig 5B). These observations mirror those seen during ZIKV infection of late term organoids and in postnatal mice (95, 99, 108). Other ZIKV isolates, including PRV, did not replicate in E19 brain slice cultures (Fig. 5B).

**Figure 5.**
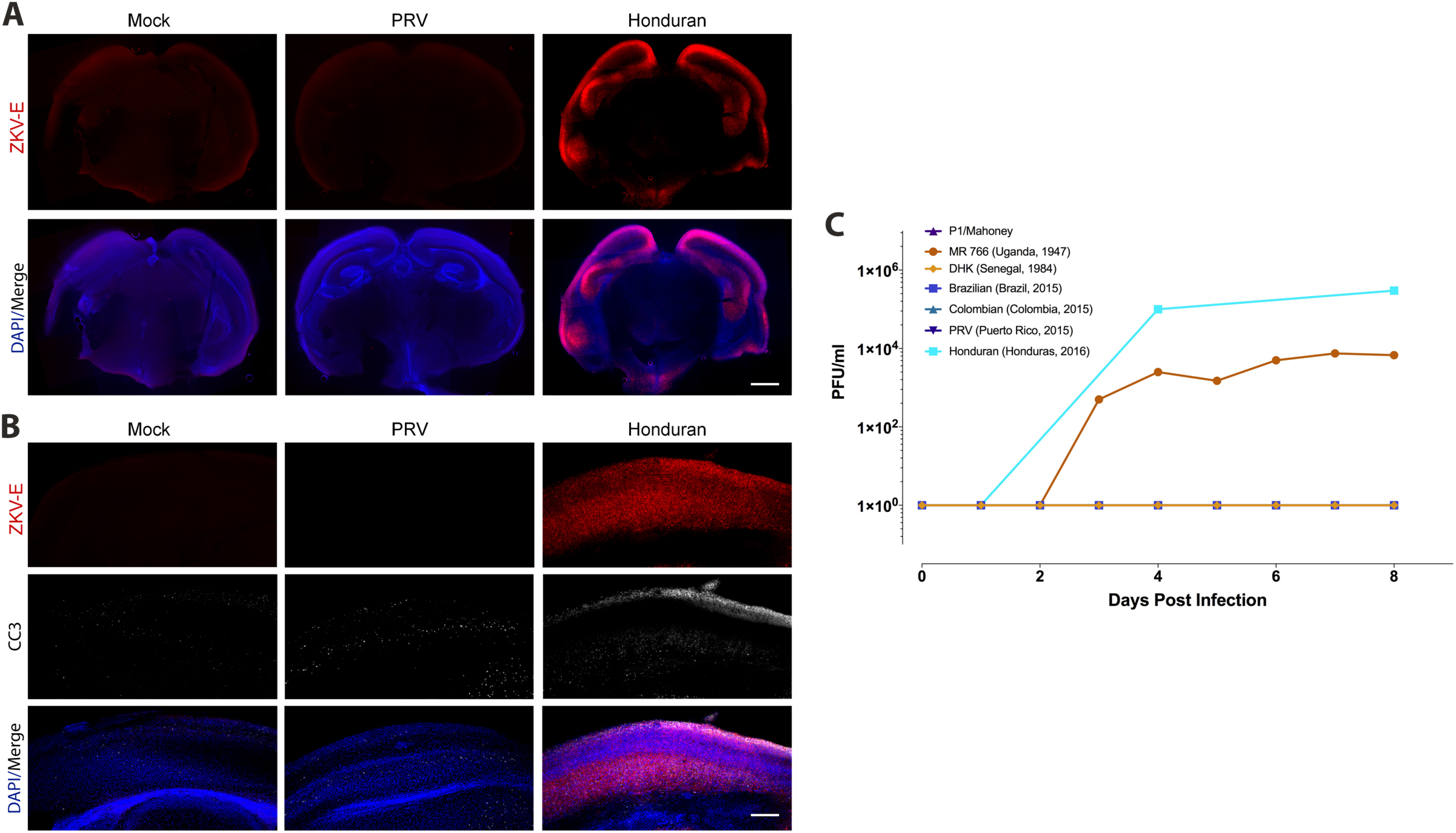
Replication of two ZIKV isolates in late gestation neural tissue. **(A, B)** Brain slice cultures from E19 embryos were infected with 10^5^ pfu of the indicated ZIKV isolates, and at 8 dpi were fixed and stained with pan-flavivirus antibody against the E glycoprotein (ZIKV-E), cleaved caspase 3 (CC3) and DAPI. Scale bar is 500 and 250 µm in **(A)** and **(B)**. **(C)** Time course of replication of multiple isolates of ZIKV in E19 brain slice cultures. P1/Mahoney, DHK, Colombian, and PRV isolates did not replicate are the lines are superimposed and not readily visible.

### Effects of ZIKV infection on neocortical layer formation

The spectrum of neuronal malformations associated with clinical cZVS is no longer limited to microcephaly. It includes brain calcifications, lissencephaly, ventriculomegaly, cerebellar hypoplasia, and brainstem dysfunction, leading to clinical symptoms including seizures and spasticity (4, 5, 8, 11-15). The wide spectrum of neuropathology seen in patients with cZVS suggests that multiple aspects of brain development may be affected, including neuronal migration. To gain insight as to whether ZIKV replication may alter this behavior, GFP-encoding DNA was electroporated *ex utero* into the radial glial progenitor cells (109) that line the ventricular surface of E15 brains prior to generation of the organotypic cultures, and infected with ZIKV 24 hr later. IF of organotypic cultures from these brains revealed that ZIKV infection impeded neuronal redistribution as judged by a lower percentage of electroporated cells reaching the most superficial layers of the neocortex 4 dpi (Fig. 6B, C). All of the ZIKV isolates tested impaired neuronal colonization into the cortical plate (Fig. 6A-C).

**Figure 6.**
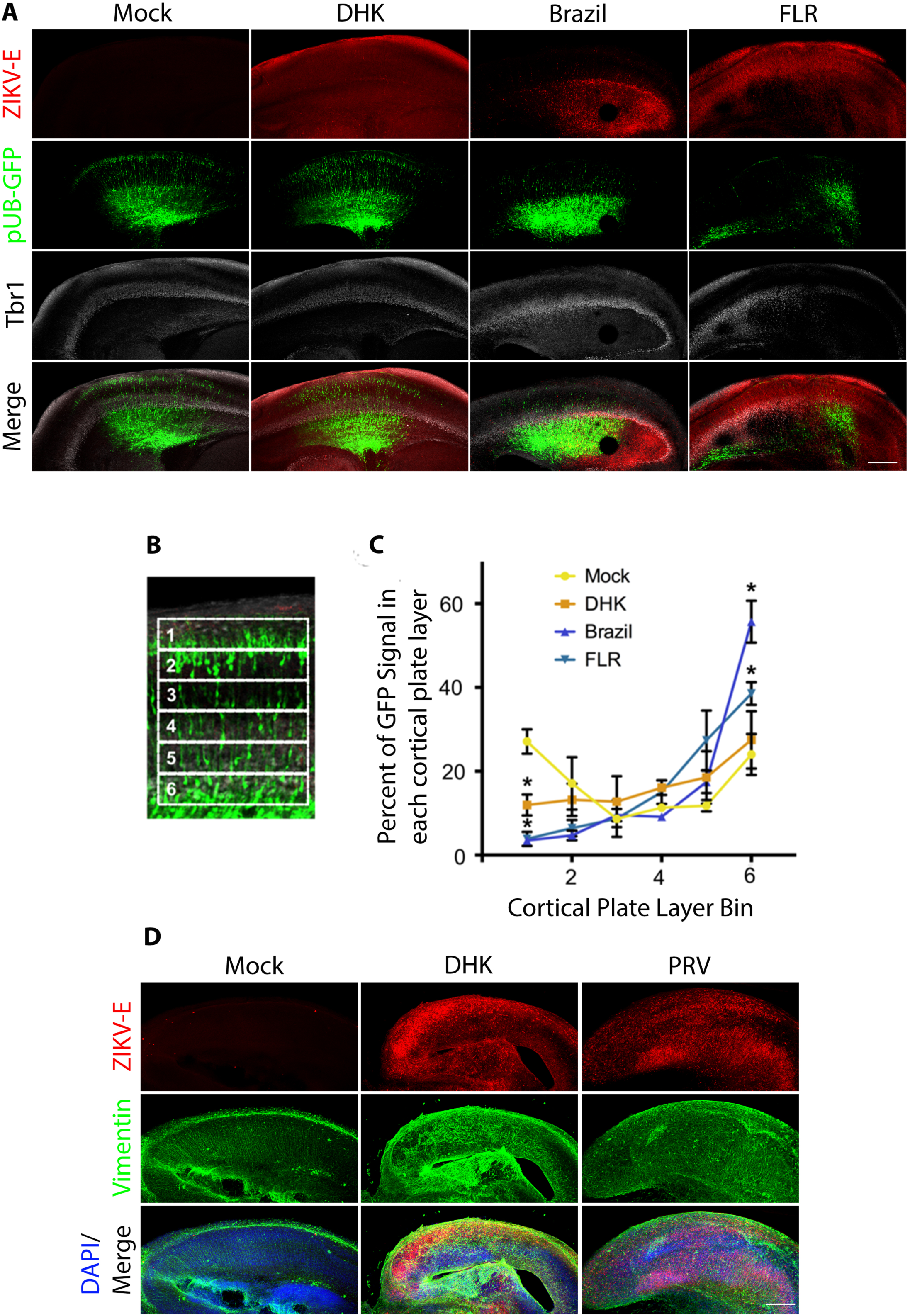
Neuronal migration is impaired during ZIKV infection. **(A)** Plasmid encoding GFP DNA was electroporated into E19 brains ex utero prior to production of slice cultures. Twenty-four hours later, the slice cultures were infected with 10^5^ pfu of the indicated ZIKV isolates, and at 4 dpi were fixed and stained with pan-flavivirus antibody against the E glycoprotein (ZIKV-E), antibody against cortical layer 2-4 marker Tbr1, and DAPI. **(B)** The cortical plate was divided into six regions of equal size, and the percentage of GFP fluorescent intensity in each sector over the whole six sectors was calculated. **(C)** Distribution of GFP positive cells within each sector for mock and ZIKV infected slices. 1 is the pial surface, and 6 is the ventricular surface. Data represented as mean ± SEM for each condition in each bin in **(C)**. Unpaired t-test was used to determine significance (* for p< 0.05, n=3 embryonic brains across different replicate experiments). **(D)** Sections from panel **(A)** were stained with antibody against the E glycoprotein (ZIKV-E), and antibody against vimentin to mark the radial glia progenitor (RGP) basal processes, which are the tracks upon which bipolar neurons migrate, and with DAPI. Scale bar is 250 µm for **(A)** and **(D)**.

Microscopic analysis of the infected organotypic slices also revealed that ZIKV infection perturbs the basal projections of the radial glial progenitor cells, marked by vimentin staining (Fig. 6D). Intact radial glial fibers (Fig. 6D, Mock panel) are necessary for proper neuronal migration into the cortical plate. These data support the hypothesis that ZIKV infection can cause structural changes to the basal process of the radial glial progenitor cells and, thereby, impair neuronal migration. A similar scaffold disorganization was observed in ZIKV infected organotypic human fetal brain slices (110) and is an important contributor to other brain developmental diseases (111). Impaired colonization of the cortex with postmitotic neurons is a known cause of autosomal recessive microcephaly and other genetic neurodevelopmental disorders (32). Reduced cortical density may be one mechanism by which congenital Zika virus syndrome develops.

## Discussion

Congenital ZIKV syndrome is a significant public health challenge. Microcephalic children may be born to both symptomatic or asymptomatic infected mothers, complicating proper diagnosis and treatment. ZIKV associated neuronal dysfunction may develop postnatally, and there remains a lack of understanding of the long time cognitive and physical disabilities associated with fetal ZKV syndrome (105, 107, 112-117). Many models of ZIKV infection have been established but none have or can systematically examine the consequences of viral infection across pre- and postnatal brain development (65-88, 102, 110, 118).

### Mouse brain slices as a model system

The results reported here demonstrate that organotypic brain slice cultures are ideal for studying ZIKV neurotropism. Infection of mouse embryonic brain slice cultures with ZIKV lead to identification of the sites of infection, and visualization of both cell death and impaired neuronal migration into the cortical plate. Although production of organotypic brain slices severs some axonal connections, these cultures nevertheless maintain many aspects of *in vivo* brain biology, including functional local synaptic circuitry with preserved brain architecture, vascularization, and immune composition, and can remain viable for at least 8 days in culture. They can be generated from the exact stage of embryonic development desired. Organoid cultures are unable to faithfully mimic the cellular and structural complexity of the brain, including the presence of a cortical plate, and are heterogeneous with respect to cell type composition and structure, which influences the reproducibility of any findings (91).

### Was ZIKV always neurotropic?

Two hypotheses have been suggested to explain the recent epidemic emergence of ZIKV and previously unrecognized neurotropism and congenital disease (97). One possibility is that mutations in the viral genome were selected that enhance transmission and expand tropism of the virus. Alternatively, the virus might have been randomly introduced into large immunologically naïve populations, and the increased numbers of cases revealed previously undetected syndromes. All ZIKV isolates examined, from 1947 to the present, replicated in early and mid-gestation embryonic brain slice cultures (E13 and E15, respectively), demonstrating that ZIKV neurotropism is not a newly acquired characteristic. This observation suggests that the earliest human ZIKV infections could have led to neuronal pathology that was too rare for a clear association to be established, especially in areas with poor health infrastructure. Our findings do not rule out the possibility that mutations within the ZIKV genome have been selected that enhance viral neuroinvasion.

### Cell type and developmental stage specificity

The greatest risks for developing cZVS are associated with ZIKV infection during the first and second trimester (13, 58). Our results show that two ZIKV isolates can replicate in brain slice cultures generated from E19 embryos, equivalent to the third trimester of human gestation, yet not all ZIKV isolates replicated in E19 brain slice cultures. For example, the PRV isolate, which replicated in E13 and E15 brain slice cultures, did not replicate in E19 cultures. This observation suggests that the changing cellular environment during embryonic development influences viral neurotropism. Fewer than ten amino acids distinguish PRV from MR766 and the Honduran isolate, but which is the genetic determinant of replication in E19 brain slice cultures will require further study. Recently it was shown that the PRV isolate replicated in the brain of 1 day old mice inoculated subcutaneously (119). The discrepancy with our results is unexplained, but could involve the genetic background of the mouse or the virus.

Additional brain developmental deficits associated with cZVS have been identified since the first reports of the association of ZIKV infection with microcephaly. These observations are consistent with our understanding that a decrease in neuron number due to either inhibition of RGP proliferation or apoptosis reduces brain size. When apoptosis predominates, areas of programmed cell death contribute not only to an overall microcephalic phenotype, but facilitate the development of neocortical defects such as intracerebral calcifications. Additionally, the neuronal migration defects observed in areas of ZIKV-envelope staining are more typical of the neuropathology associated with lissencephaly. Together these data suggest that multiple mechanisms may contribute to the neuropathology in children diagnosed with cZVS.

## Materials and Methods

### Ethics Statement

All experiments were performed in accordance with guidelines from the Guide for the Care and Use of Laboratory Animals of the NIH. Protocols were reviewed and approved by the Institutional Animal Care and Use Committee (IACUC) at Columbia University School of Medicine (assurance number AC-AAAR5408).

### Cell and organotypic brain slice cultures from embryonic mice and viruses

Vero cells were grown in Dulbecco’s modified Eagle medium (Invitrogen, Carlsbad, CA), 10% fetal calf serum (HyClone, Logan, UT), and 1% penicillin-streptomycin (Invitrogen).

Timed pregnant Swiss Webster mice (E13, E15 and E19) were purchased from Taconic Labs; E1 was defined as the day of confirmation of sperm-positive vaginal plug. Mice were sacrificed and fetuses were harvested. Fetal brains were dissected into ice-cold artificial cerebrospinal fluid (ACSF) consisting of 125mM NaCl, 5mM KCl, 1.25mM NaH_2_ PO_4_, 1mM MgSO_4_, 2mM CaCl_2_, 25mM NaHCO_3_, and 20mM glucose, pH7.4, 310 mOsm1^-1^. Brains were embedded in 4% low melting point agarose dissolved in ACSF and sliced into 300µm coronal sections using a vibratome (Zeiss). Slices were maintained on 0.4 µm, 30mm diameter Millicell-CM inserts (Millipore) in cortical culture medium (CCM) containing 25% Hanks balanced salt solution, 47% basal MEM, 25% normal horse serum, 1X penicillin-streptomycin-glutamine, and 30% glucose. Cultures were maintained in a humidified incubator at 37°C with constant 5% CO_2_ supply. Zika viruses (ZIKV) MR766 (Ugandan origin), DAK (Senegal), FLR (Colombia), PRVABC59 (Puerto Rico) and R103451 (Honduras) were obtained from BEI Resources. Brazilian isolate (ZIKV-Paraiba/2015) was kindly provided by Lucia Gama (Johns Hopkins School of Medicine, Baltimore Maryland). All viruses were propagated and assayed in Vero cells. Viral titers were determined by plaque assay.

### *Ex-utero* electroporation

Plasmids encoding GFP were transfected by intraventricular injection into the ventricle of dissected embryonic brains at the appropriate time. DNA was mixed with colored non-toxic dye and 1µg of nucleic acid was injected into the ventricular space using a high gauge needle made from glass capillary tube. Post injection, five pulses of electrical current (50V, 5ms each, with 1s intervals) were applied by directly placing electrodes on the outer surface of the brain, angled along the lateral aspect of the neocortex adjacent to the lateral ventricle targeted by injection. Brain slices were generated following transfection and cultures were maintained up to 8 days post electroporation.

### Indirect immunofluorescence microscopy

Either 72 h, 96h or 8 days post infection, the medium was removed from Zika virus infected or uninfected organotypic brain slices cultures and the cultures were placed overnight in 4% paraformaldehyde (PFA) fixative dissolved in 1X PBS at 4 °C. Following fixation, cultures were incubated in blocking solution of PBS, 0.3% Triton X-100 and 3% donkey serum. Cultures were incubated overnight in at 4 °C in blocking solution containing appropriate primary antibodies. Sections were washed in 1X PBS, and incubated in the presence of fluorophore-conjugated secondary in blocking solution. Sections were mounted on slides using Aqua-Poly/Mount (Polysciences, Inc) and imaged using a iZ80 laser scanning confocal microscope (Olympus FV100 spectral confocal system).

### Plaque assay

Vero cells were seeded on 60mm plates for approximately 70% confluence at the time of plaque assay. Next, 100λ portions of serial 10-fold virus dilutions were incubated with cells for 1 h at 37 °C. Two overlays were added to the infected cells. The first overlay consisted of 2 ml of 1 DMEM, 0.8% Noble agar, 0.1% bovine serum albumin, 40 mM MgCl_2_, and 10% bovine calf serum. After solidification, a second liquid overlay was added that was composed of 1 DMEM, 0.1% bovine serum albumin, 40 mM MgCl_2_, 0.2% glucose, 2 mM pyruvate, 4 mM glutamine, and 4 mM oxaloacetic acid. The cells were incubated at 37 °C for 6-8 days and developed by using 10% trichloroacetic acid and crystal violet.

### Virus infections

Organotypic brain slice cultures were infected with 10^5^ pfu of ZIKV isolates: MR766 African, DHK Senegal, Brazilian, FLR, PRVABC59, and R103451 Honduras, or poliovirus (P1/Mahoney). Virus was allowed to adsorb to the slices for 1h at 37 °C. The inoculum was removed, and the slices were washed 2X in 1X PBS. Infected slices were cultured in CCM for 8 days. 500µl of culture medium was removed and replenished at the described points post infection.

### Antibodies

Antibodies used in this study were chicken polyclonal against vimentin (Millipore, AB5733, 1:1000 dilution), pan-flavivirus E glycoprotein (Millipore, MAB 10216), rabbit polyclonal against CDP (Santa-Cruz, SC-1302), and cleaved caspase 3 (Cell Signaling) Donkey fluorophore-conjugated secondary antibodies (Jackson Labs, 1:500 dilution) were used together with DAPI (4’,6-diamidino-2-phenylindole, Thermo Scientific, 62248, 1:1,000 dilution).

### Data Analysis

GraphPad Prism software was used to analyze all data. Log10-transformed titers were used for graphing the results of plaque assays.

## Acknowledgments

This project was supported by NIH GM105536 to R.B.V., NINDS F30NS095577 to D.J.D., and NIH AI121944 and AI102597 to V.R.R.

We thank the members of the Vallee lab, Drs. Alex Baffett, Hynek Wichterle, Franck Polleux, Gregg Gundersen, Yosef Sabo, Elisa Canepa, Stav Kameny, and Leslie Vosshall for technical expertise and feedback, and Drs. Connie Cepko and Lucia Gama for providing reagents.

---

**Supplemental Figure 1.**
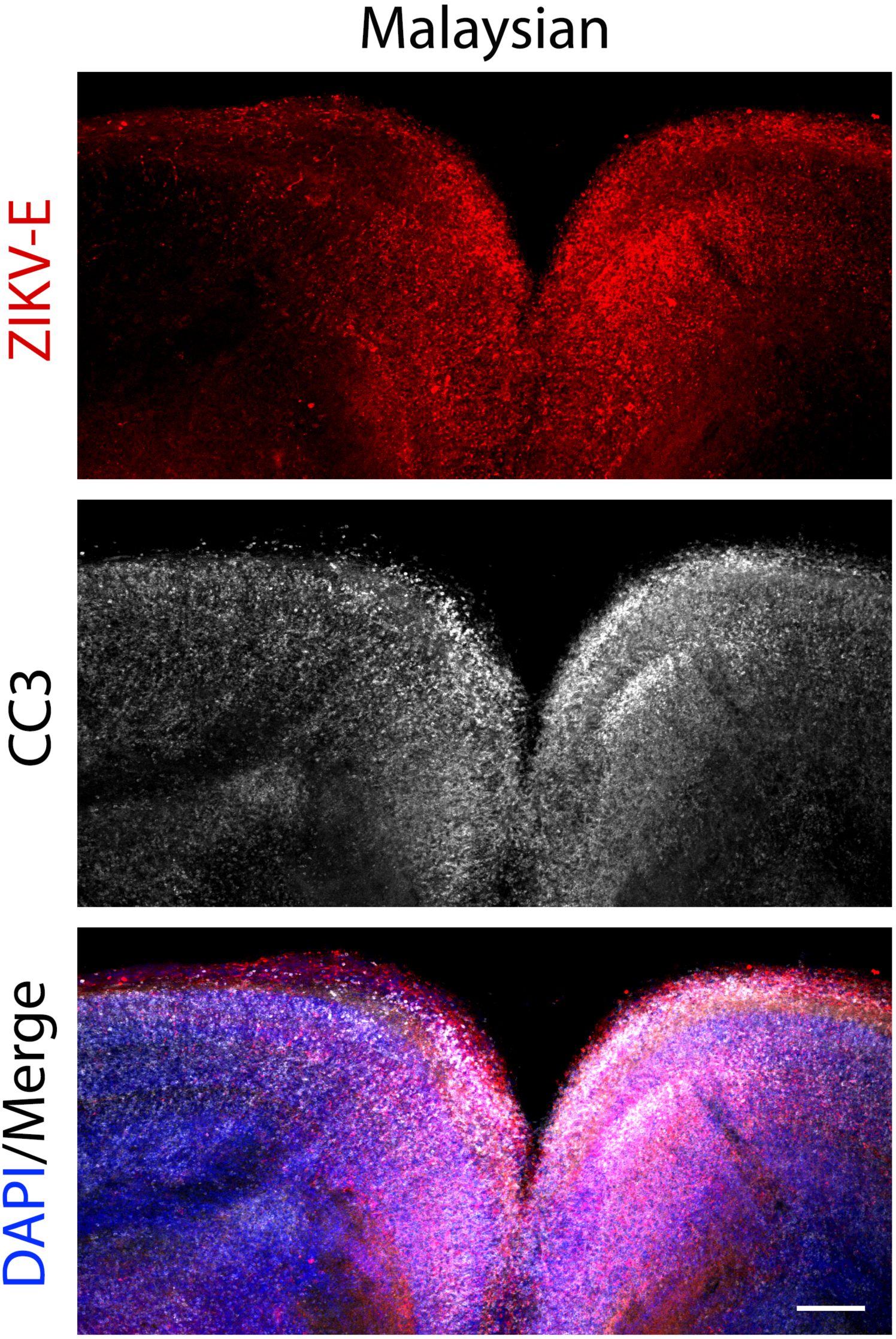
Tissue distribution and replication of Malaysian ZIKV isolate in E15 organotypic mouse brain slices. Brain slice cultures from E15 embryos were infected with 10^5^ pfu of the Malaysian ZIKV isolate, and at 4 dpi were fixed and stained with pan-flavivirus antibody against the E glycoprotein (ZIKV-E), cleaved caspase 3 (CC3) and DAPI. Scale bar is 500 µm.

